# Human intestinal enteroids with inducible neurogenin-3 expression as a novel model of gut hormone secretion

**DOI:** 10.1101/579698

**Authors:** Alexandra L. Chang-Graham, Heather A. Danhof, Melinda A. Engevik, Catherine Tomaro-Duchesneau, Umesh C. Karandikar, Mary K. Estes, James Versalovic, Robert A. Britton, Joseph M. Hyser

## Abstract

**Background:** Enteroendocrine cells (EECs) are specialized epithelial cells that produce molecules vital for intestinal homeostasis, but due to their limited numbers, in-depth functional studies have remained challenging. Human intestinal enteroids (HIEs) that are derived from intestinal crypt stem cells are a biologically relevant *in vitro* model of the intestinal epithelium. HIEs contain all intestinal epithelial cell types; however, like the intestine, HIEs spontaneously produce few EECs, which limits their study.

**Methods:** To increase the number of EECs in HIEs, we used lentivirus transduction to stably engineer jejunal HIEs with doxycycline-inducible expression of neurogenin-3 (*NGN3*), a transcription factor that drives EEC differentiation (tet*NGN3*-HIEs). We examined the impact of *NGN3* induction on EECs by quantifying the increase in the enterochromaffin cells and other EEC subtypes. We functionally assessed secretion of serotonin and EEC hormones in response to norepinephrine and rotavirus infection.

**Results:** Treating tet*NGN3*-HIEs with doxycycline induced a dose-dependent increase of chromogranin A (ChgA)-positive and serotonin-positive cells, demonstrating increased enterochromaffin cell differentiation. Despite increased ChgA-positive cells, other differentiated cell types of the epithelium remained largely unchanged by gene expression and immunostaining. RNA sequencing of doxycycline-induced tet*NGN3*- HIEs identified increased expression of key hormones and enzymes associated with several other EEC subtypes. Doxycycline-induced tet*NGN3*-HIEs secreted serotonin, monocyte chemoattractant protein-1, glucose-dependent insulinotropic peptide, peptide YY, and ghrelin in response to norepinephrine and rotavirus infection, further supporting the presence of multiple EEC types.

**Conclusions:** We have combined HIEs and inducible-*NGN3* expression to establish a flexible *in vitro* model system for functional studies of EECs in enteroids and advance the molecular and physiological investigation of EECs.

**Synopsis:** Enteroendocrine cells have low abundance but exert widespread effects on gastrointestinal physiology. We engineered human intestinal enteroids with inducible expression of neurogenin-3, resulting in increased enteroendocrine cells and facilitating investigations of host responses to the dynamic intestinal environment.

## Introduction

The gastrointestinal (GI) epithelium is the largest sensory interface between host and environment and must both detect and communicate luminal contents to the host [1]. The GI lumen is a complex mixture of dietary nutrients and their breakdown products, microorganisms and their metabolites, as well as irritants, toxins, and drugs. Microorganisms in particular create a dynamic ecosystem in the GI lumen with both beneficial and detrimental effects. For example, the gut microbiome is important for degradation of complex carbohydrates and polysaccharides into short chain fatty acids (SCFAs) such as acetate, butyrate, and propionate that can be used as nutrients by the host or other microbes. SCFAs also modulate a variety of important physiological effects such as inflammation and gut motility [2]. By contrast, pathogenic microorganisms invade the epithelium or produce toxins that threaten the GI epithelial barrier and thus necessitate activation of host defenses. Therefore, integrating signals from luminal stimuli to the host for coordinated and appropriate physiological responses is complex, which has complicated efforts to study the molecular mechanisms governing these processes.

Microbiome-to-epithelium communication is multifaceted and includes receptors on various cell types, including enterocytes, goblet cells, Paneth cells and stem cells. Recent evidence shows that enteroendocrine cells are one of the most important mediators of communication between the GI lumen and host [3]. Enteroendocrine cells (EECs) are a rare cell lineage found throughout the length of the GI tract (<1% of GI epithelial cells), yet they have a significant impact on human physiology [4-6]. In response to host, microbial, and environmental stimuli, subtypes of EECs synthesize and secrete hormones including gastrin (G cells), somatostatin (D cells), enteroglucagon/peptide YY (PYY) and glucagon-like peptide 1 (GLP-1) (L cells), neurotensin (N cells), pro-γ-melanocyte-stimulating hormone (MSH) cells, and serotonin (enterochromaffin cells) [7-9]. The broad array of enteroendocrine cell types reflects the diversity of critical physiological functions including the coordination of both local and systemic responses by the endocrine and nervous systems to the stimuli present in the lumen. Enterochromaffin cells are the most common EEC subtype, comprising ∼40% of the EECs. These cells synthesize and secrete the neurotransmitter serotonin in response to various physiological stimuli, including microbial metabolites, irritants, toxins, and infection [10-12]. Given the importance of serotonin in mammalian physiology and greater abundance among EECs, many studies focus on enterochromaffin cells, but due in part to their rarity, how other EEC subtypes function in the context of the gut environment remains less well studied.

Studying the interaction of EECs and stimuli in the GI lumen has been challenging due to the limited numbers of these cells in the intestinal epithelium and the lack of appropriate non-cancer derived *in vitro* models. Generation of enteroids is a recent technology that has revolutionized the study of intestinal epithelial cells. They have been generated from several mammalian hosts, including mouse and human, and recapitulate the GI physiology of the donor species [13-16]. This complex epithelial culture system is derived from adult stem cells isolated from intestinal tissue or biopsy samples that can be maintained and expanded in culture as an *in vitro* model system [13-15, 17-21]. Thus, enteroids offer a promising new tool to study non-transformed, non-cancerous human EECs in an *in vitro* culture system [17].

Previous studies have established that neurogenin 3 (Ngn3) is a key transcription factor for EEC differentiation [22-36]. Consistent with the role of Ngn3 in EEC fate, overexpression of *NGN3* in multiple systems has been shown to increase EEC numbers [37, 38]. *In vitro*, adenoviral-based *NGN3* overexpression in neonatal mouse jejunal intestinal spheres induced a threefold increase in the number of chromogranin A (ChgA)–positive EECs [39]. In human intestinal organoids (HIOs) derived from pluripotent stem cells, overexpression of *NGN3* by an adenoviral vector or tetracycline-inducible lentiviral vector also increased ChgA-positive EECs [29, 40]. In this study, we generated a new model system using jejunal human intestinal enteroids (HIEs) engineered to overexpress *NGN3* from a tetracycline-inducible promoter (tet*NGN3*) to drive EEC differentiation. These tet*NGN3*-HIEs exhibit a doxycycline dose-dependent increase in ChgA expression in both 3-dimensional culture and as 2-dimensional monolayers, as well as upregulation of markers for multiple EEC subtypes, including enterochromaffin cells, L cells and K cells. In response to microbial and host stimuli, these induced EECs secreted serotonin, monocyte chemoattractant protein-1 (MCP-1), glucose-dependent insulinotropic peptide (GIP), peptide YY (PYY), and ghrelin. Thus, tet*NGN3*-HIEs are a new and physiologically relevant model system to study EEC-based communication pathways in response to both microbial and host stimuli.

## Results

### Creation and propagation of inducible tetNGN3-HIEs

Since host-microbe interactions via EECs has a profound impact on host physiology, we sought to develop a new model for screening, identification, and testing of microbe-induced EEC responses. Due to the low abundance of EECs within intestinal tissue and human intestinal enteroids (HIEs), we aimed to increase EEC abundance through inducible overexpression of *NGN3* using lentivirus transduction to introduce a doxycycline-inducible NGN3 expression cassette into an HIE [72, 73]. In preparation for transduction, jejunal HIEs were grown high Wnt complete media with growth factors (hW-CMGF+) to enrich the stem cell population, which was evidenced by the majority of HIEs exhibiting a cystic morphology with multiple small buds (**Fig. 1A, left panel**). Growth in high Wnt media was important for increasing the success rate of the transduction. After lentivirus transduction, geneticin selection initially increased sloughing of dead cells from the HIEs, and full selection of transduced tet*NGN3*-HIEs occurred after ∼5 weeks (**Fig. 1A, right panel**).

**Figure 1:**
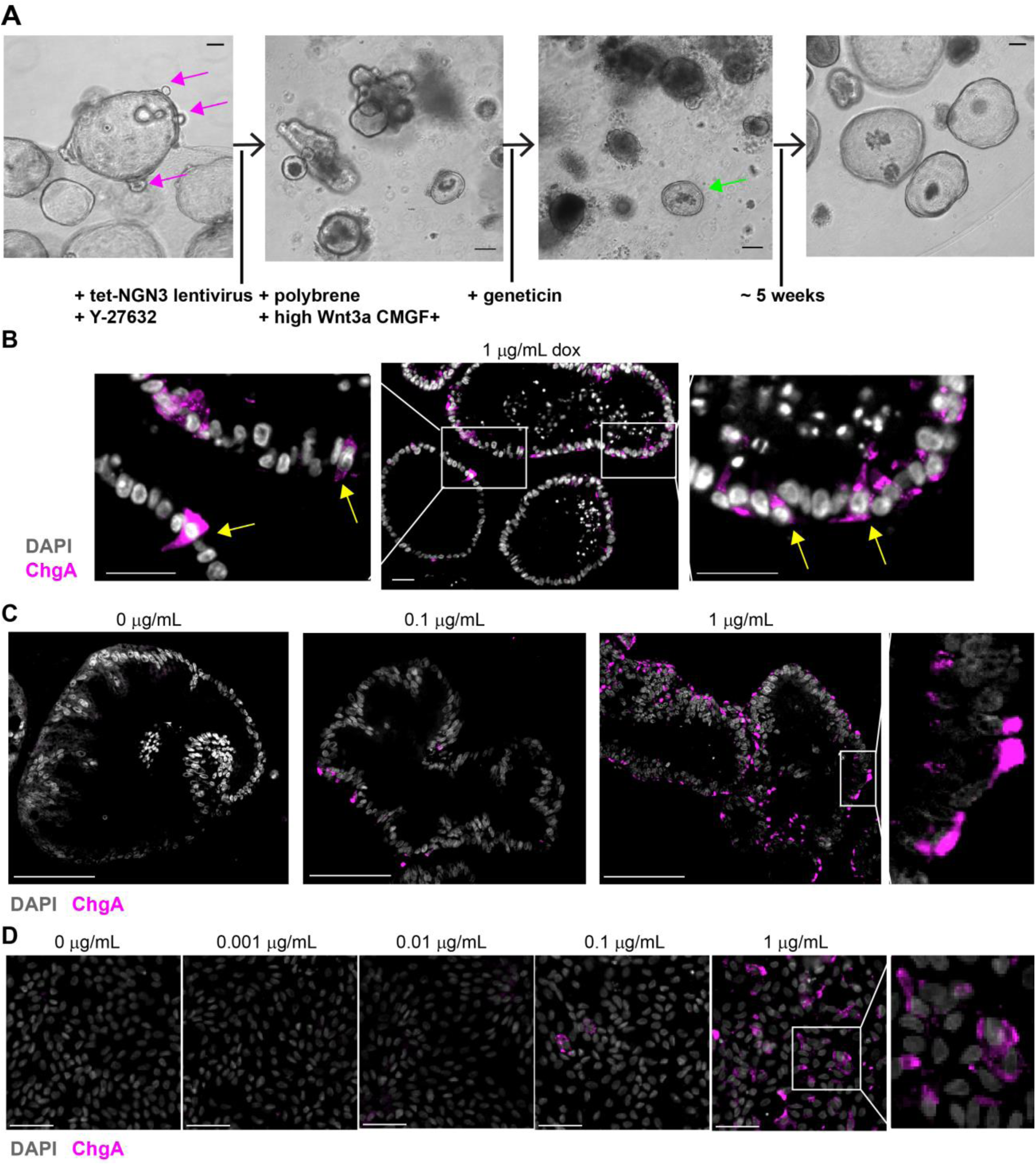
**(A)** Production pipeline for creating tet*NGN3*-HIEs using lentivirus transduction. Jejunum HIEs grown in high Wnt CMGF+ media increases stem cells, evidenced by cysts and crypt buds (pink arrows), followed by inoculation with lentivirus. Reseeding the HIEs in Matrigel is followed by geneticin selection for approximately 5 weeks, so that only HIEs with tet*NGN3* construct survive (green arrow) (scale bar = 100µm). Images were acquired using a 10X Plan Fluor (NA 0.3) phase contrast objective on an inverted Nikon TiE microscope with an ORCA-Flash 4.0 sCMOS camera and Nikon Elements software. **(B, C, D)** Doxycycline induces enterochromaffin marker chromogranin A (ChgA) expression in tet*NGN3*-HIEs. Tet*NGN3*-HIEs were fixed and stained for ChgA (Alexa Fluor 488 [pink]) to mark enterochromaffin cells and counterstained with DAPI (gray). **(B)** 3D tet*NGN3*-HIEs were differentiated for 4 days with 1 µg/mL doxycycline (scale bars = 25 µm). ChgA-positive cells (inset) were observed (yellow arrows). **(C, D)** tet*NGN3*-HIEs as **(C)** 3D cultures and **(D)** flat monolayers were differentiated for 4 days with 0, 0.1, and 1 µg/mL doxycycline and also 0.01 and 0.001 µg/mL doxycycline in flat monolayers. 3D culture images were acquired using a 20X Plan Apo (NA 0.75) DIC objective on an upright Nikon Eclipse 90i microscope with a Photometrics CoolSNAP HQ2 camera and Nikon Elements software (scale bar = 100 µm). Flat monolayer images were acquired using a 20X Plan Apo (NA 0.75) DIC objective on an inverted Nikon TiE microscope with an ORCA-Flash 4.0 sCMOS camera and Nikon Elements software (scale bar = 50 µm).

To confirm the ability of *NGN3* to drive EEC differentiation we used immunofluorescence (IF) staining for chromogranin A (ChgA) as a marker of endocrine cells, which are often used to assess increases in EEC numbers upon *NGN3* overexpression [29, 39, 40]. First, the number of ChgA-positive cells present in the tet*NGN3*-HIEs after induction with 1 µg/mL doxycycline was assessed (**Fig. 1B**). In paraffin-embedded slices of 3D tet*NGN3*-HIE cultures, we observed abundant ChgA-positive cells in both apical and basolateral areas of cells (**Fig 1B**). In contrast to the doxycycline-induced tet*NGN3*-HIEs, very few ChgA-positive cells were observed in the absence of doxycycline (**Fig. 1C, left panel**). However, induction of tet*NGN3* increased the number of ChgA-positive cells in a doxycycline dose-dependent manner (**Fig. 1C**). A similar doxycycline dose-dependent increase in ChgA-positive cells was observed in two-dimensional (2D) flat monolayers (**Fig. 1D**). Of note, the ChgA-positive cells in both 3D and 2D formats exhibited the polygonal cell shape typically associated with enterochromaffin cells [41].

Using image analysis software, we quantified the abundance of ChgA-positive cells from the different doxycycline-induction conditions above. In the absence of doxycycline, both 3D and monolayer tet*NGN3*-HIEs exhibited few to no ChgA-positive cells. In contrast, addition of 0.1 µg/mL and 1 µg/mL doxycycline to 3D tet*NGN3*-HIEs resulted in a dose-dependent increase in the number of ChgA-positive cells (p<0.0001) (**Fig. 2A, left panel**). This pattern was also observed in 2D flat tet*NGN3*-HIEs monolayers, with doxycycline concentrations at or below 0.01 µg/mL exhibiting ∼0.4% ChgA-positive cells, similar to the non-induced tet*NGN3*-HIEs (**Fig. 2A, right panel**). Treatment of tet*NGN3*-HIEs with 0.1 µg/mL doxycycline resulted in ∼5% ChgA-positive cells, about a 12-fold increase, while addition of 1 µg/mL doxycycline resulted in ∼40% ChgA-positive cells (**Fig. 2A right panel**). Thus, induction of tet*NGN3* correlated with an increase in ChgA-positive cells, supporting our premise that overexpression of *NGN3* would drive EEC differentiation in HIEs. The tet*NGN3*-HIEs have stably maintained the tet*NGN3* transgene and exhibited the doxycycline-induced increase in ChgA-positive cells (detected by IF staining) for >10 months. Additionally, tet*NGN3-*HIEs maintained inducible expression after storage in liquid nitrogen (data not shown).

**Figure 2:**
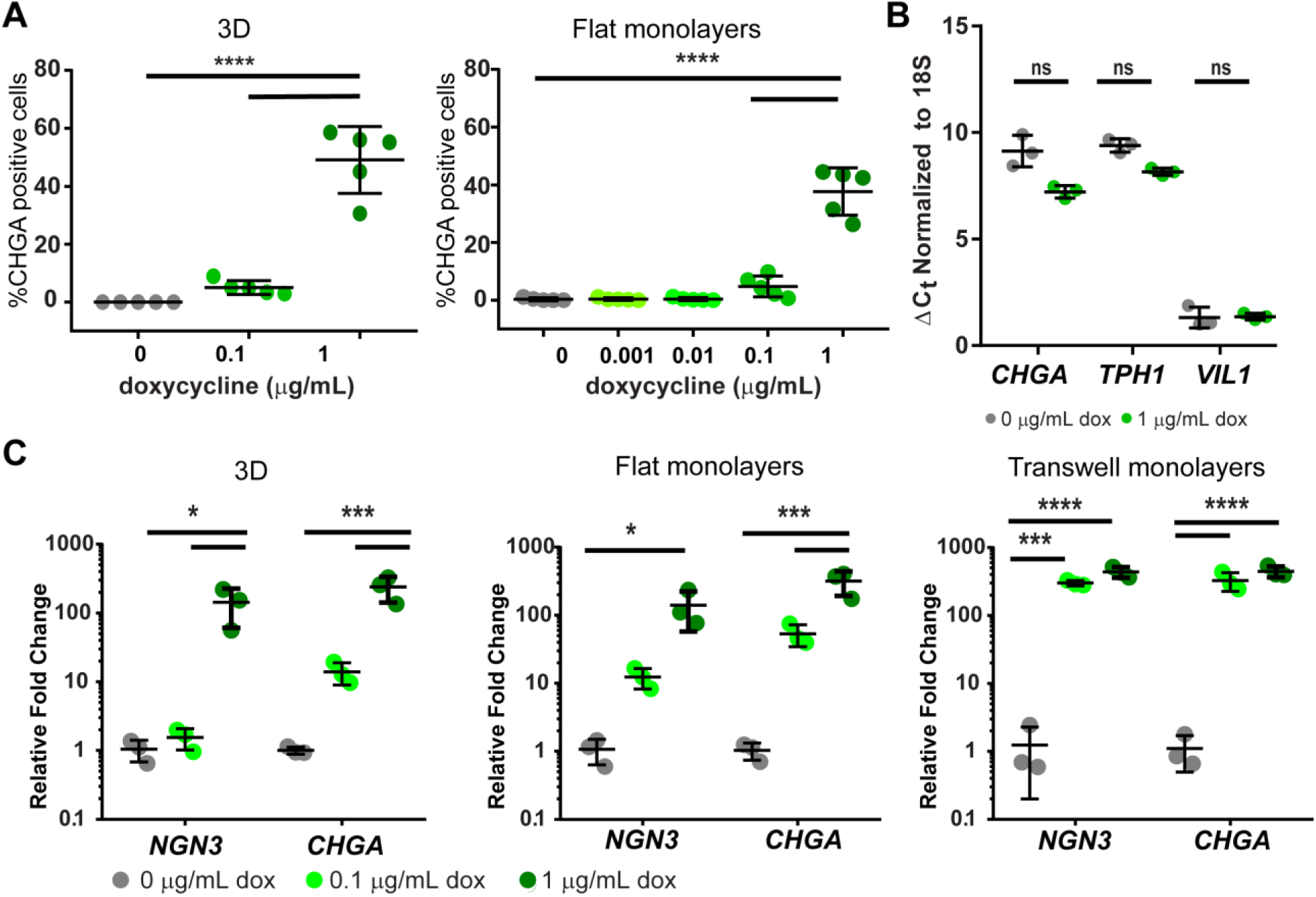
Doxycycline induces enteroendocrine cell lineage in tet*NGN3*-HIEs. **(A)** Increased doxycycline concentration increases the percentage of ChgA-positive cells in differentiated tet*NGN3*-HIE 3D cultures and flat monolayers. Images were analyzed with Nikon elements software with 5 images per condition with an average of 220 nuclei per image in 3D cultures and 1400 nuclei per image in flat monolayers. (n = 2 biological replicates) **(B)** qPCR of *CHGA (*chromogranin A), *TPH1* (tryptophan hydrolase-1), and *VIL1* (villin) mRNA transcripts of parental jejunum 3D enteroids were treated with 0 or 1 µg/mL doxycycline and normalized to 18S mRNA. (n = 3 biological replicates) **(C)** qPCR of *NGN3* and *CHGA* mRNA transcripts normalized to 18S mRNA transcripts in tet*NGN3* 3D cultures, flat monolayers, and transwell monolayers. (n = 3 biological replicates) (ns, not significant, *p<0.05, ***p<0.001, ****p<0.0001).

### Induction of enteroendocrine cell differentiation

To confirm that doxycycline treatment alone did not induce EEC differentiation in HIEs, we measured mRNA transcript levels after treating the parental (non-transduced) jejunum HIEs with 0 or 1 µg/mL doxycycline. We quantitated the mRNA levels of the enterochromaffin cell markers *CHGA* and *TPH1* and the enterocyte marker villin (VIL1) by quantitative polymerase chain reaction (qPCR) (**Fig. 2B**). No significant difference between treatment groups was present in any of the cell markers examined, indicating that doxycycline alone does not impact EEC differentiation in the absence of *NGN3* expression (**Fig. 2B**).

In a complementary approach to demonstrate that induction of tet*NGN3* increases EEC differentiation, we correlated the increase in *NGN3* and *CHGA* transcripts in tet*NGN3*-HIEs treated with 0, 0.1, and 1 µg/mL doxycycline, and compared gene expression in the three different HIE formats (i.e., 3D culture, flat monolayers, and transwell monolayers), summarized in **Table 3**. Overall, we observed a doxycycline dose-dependent increase in both *NGN3* and *CHGA* expression in all three formats of the tet*NGN3*-HIEs. Treatment with 1 µg/mL doxycycline significantly increased both enterochromaffin cell markers in all three culture formats (**Fig. 2C**). However, there were some notable differences in the expression profiles between each type of HIE format. Treating 3D tet*NGN3*-HIEs with 0.1 µg/mL doxycycline induced *NGN3* and *CHGA* expression to a lesser degree than 0.1 µg/mL doxycycline treatment of flat or transwell monolayers (**Fig. 2C**), which may be due to limited penetrance of doxycycline through the Matrigel used for 3D cultures. Further, transwell monolayers treated with 0.1 µg/mL doxycycline have a greater fold induction of *NGN3* and *CHGA* expression than in flat monolayers (**Fig. 2C**). This difference may be the result of greater cell surface contact with the media in transwell monolayers, which have both apical and basolateral access to the media, than in flat monolayers. Together, these results demonstrate that the inducible tet*NGN3-*HIEs are a tunable and versatile system for increased differentiation of enterochromaffin cells in each of the HIE culture formats.

**Table 3:** Fold Changes in *NGN3* and *CHGA* Expression with Doxycycline Treatment

| tetNGN3-HIE format | *NGN3* |  | *CHGA* |  |
| --- | --- | --- | --- | --- |
|  | 0.1 µg/mL dox | 1 µg/mL dox | 0.1 µg/mL dox | 1 µg/mL dox |
| 3D cultures | 1.5 | 143 | 14 | 239 |
| Flat monolayers | 12 | 141 | 54 | 316 |
| Transwell monolayers | 300 | 436 | 326 | 445 |

### Characterization of epithelial cell types present in tetNGN3-HIEs

To assess the impact of *NGN3* overexpression on HIE morphology, we treated 3D and transwell monolayer preparations of the tet*NGN3*-HIEs with 0, 0.1, and 1 µg/mL doxycycline (**Fig. 3A, B**). Hematoxylin and eosin (H&E) staining of 3D (**Fig. 3A**) and transwell monolayers (**Fig. 3B**) showed only minor alterations in morphology upon induction with 0.1 µg/mL doxycycline, primarily larger nuclei and increased cell height (**Fig. 3B, middle panel**). Strong induction with 1 µg/mL doxycycline resulted in more significant morphological changes. In the 3D tet*NGN3*-HIEs, there was a marked increase in luminal (apical) cell debris (**Fig. 3A, right panel**), and in transwell monolayers we observed larger nuclei and a discontinuous apical membrane with increased shedding of cellular material (**Fig. 3B, right panel**). Further, we examined the ultrastructure of the tet*NGN3*-HIEs using transmission electron microscopy of 3D HIEs treated for 5 days with 0 or 1 µg/mL doxycycline in differentiation media (**Fig. 3C and 3D**). Most cells observed in the 0 µg/mL doxycycline-treated tet*NGN3*-HIEs had the basolateral nuclei, apical microvilli, and brush border characteristic of enterocytes (**Fig. 3C**). As expected, in tet*NGN3*-HIEs treated with 1 µg/mL doxycycline, we observed more EECs based on greater numbers of cells with electron-dense granules, including cells open to the lumen of the 3D enteroid (**Fig. 3D**).

‘Differentiation’ of HIEs by exclusion of Wnt3a from the culture media drives maturation of the different epithelial cell types [14, 17, 21, 42, 43], so we next used RT-qPCR to determine whether *NGN3* induction altered expression of cell lineage-specific marker genes in differentiated HIEs. For this we tested markers for Paneth cells (*LYSZ*), crypt base columnar stem cells (*LGR5*), goblet cells (*MUC2)*, and enterocytes [villin (*VIL1)* and sucrase isomaltase (*SI*)] (**Table 2**). First, we observed no difference in differentiation marker expression between the parental HIE line and the uninduced tet*NGN3*-HIEs, indicating that lentivirus transduction itself did not cause alterations in HIE cell differentiation (data not shown). In 3D tet*NGN3*-HIEs (**Fig. 4A, left panel**), doxycycline induction caused no significant changes in cell lineage-specific gene expression, and while *SI* exhibited a trend for lower levels, expression of *VIL1* remained unchanged. Further, in tet*NGN3*-HIE flat monolayers the overall gene expression levels were similar to the 3D tet*NGN3*-HIEs and no significant changes in lineage-specific marker genes were observed (**Fig. 4A, right panel**).

**Table 2:**
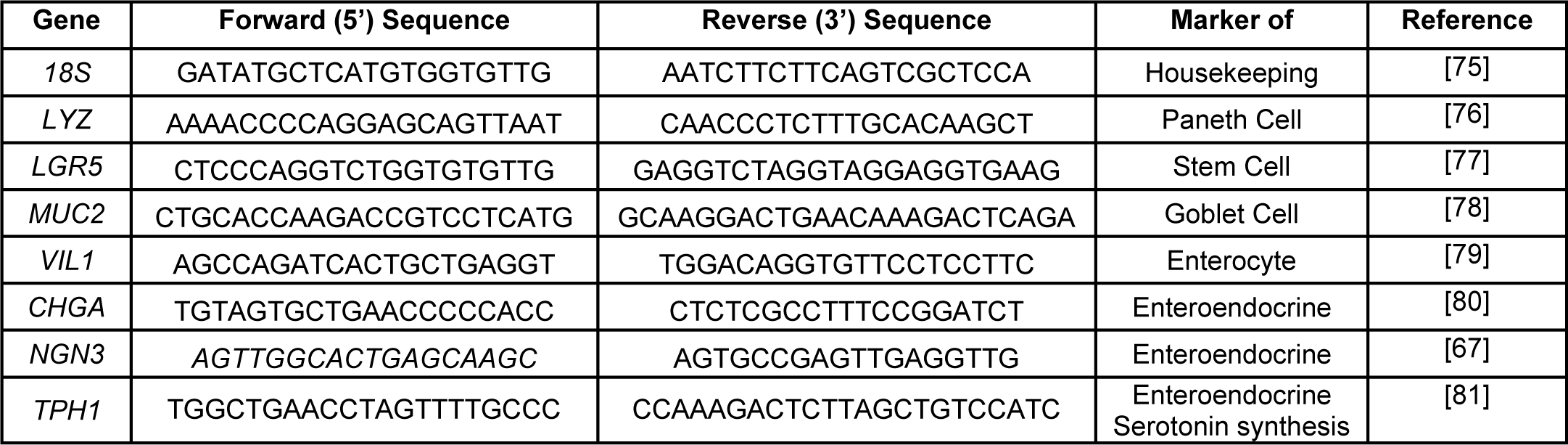
Primers used in this work

**Figure 3:**
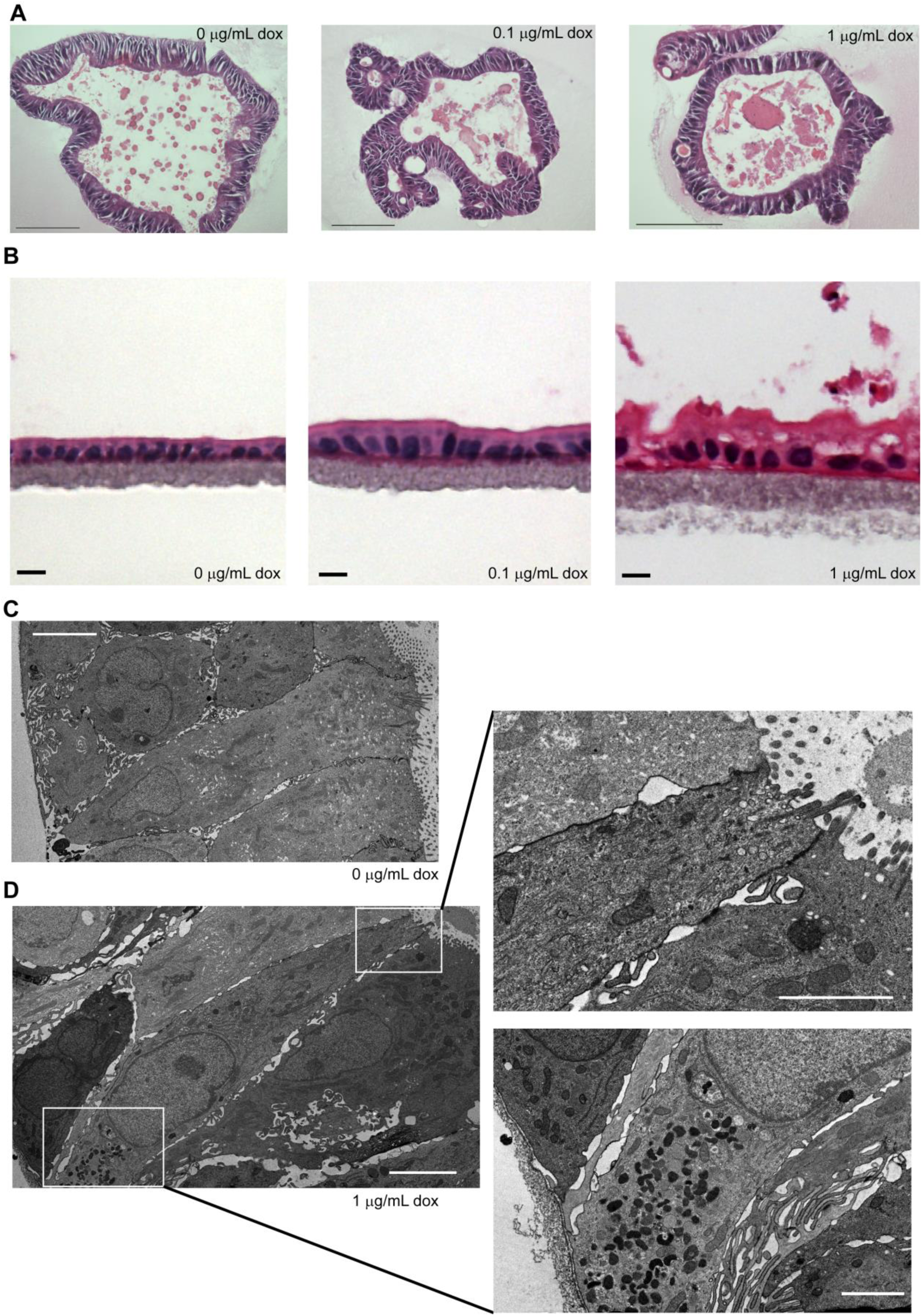
tet*NGN3*-HIEs maintain cellular morphology in culture. Hematoxylin and eosin (H&E) stained **(A)** three-dimensional (scale bar = 100µm) and **(B)** transwell monolayer (scale bar = 10µm) tet*NGN3*-HIEs treated with 0, 0.1, or 1 µg/mL doxycycline. Images were acquired using a 20X Plan Apo (NA 0.75) DIC objective on an upright Nikon Eclipse 90i microscope with a DS-Fi1-U2 camera and Nikon Elements software. Transmission electron micrographs (TEM) H&E of three-dimensional tet*NGN3*-HIEs with **(C)** 0 µg/mL (scale bar = 4µm, 1500x magnification) or **(D)** 1 µg/mL doxycycline (left scale bars = 4µm, 1500x magnification, right scale bars = 2µm, 3000x magnification). TEM sections were viewed on a Hitachi H7500 transmission electron microscope set to 80kV and collected using an AMT XR-16 digital camera and AMT Image Capture, v602.600.51software.

**Figure 4:**
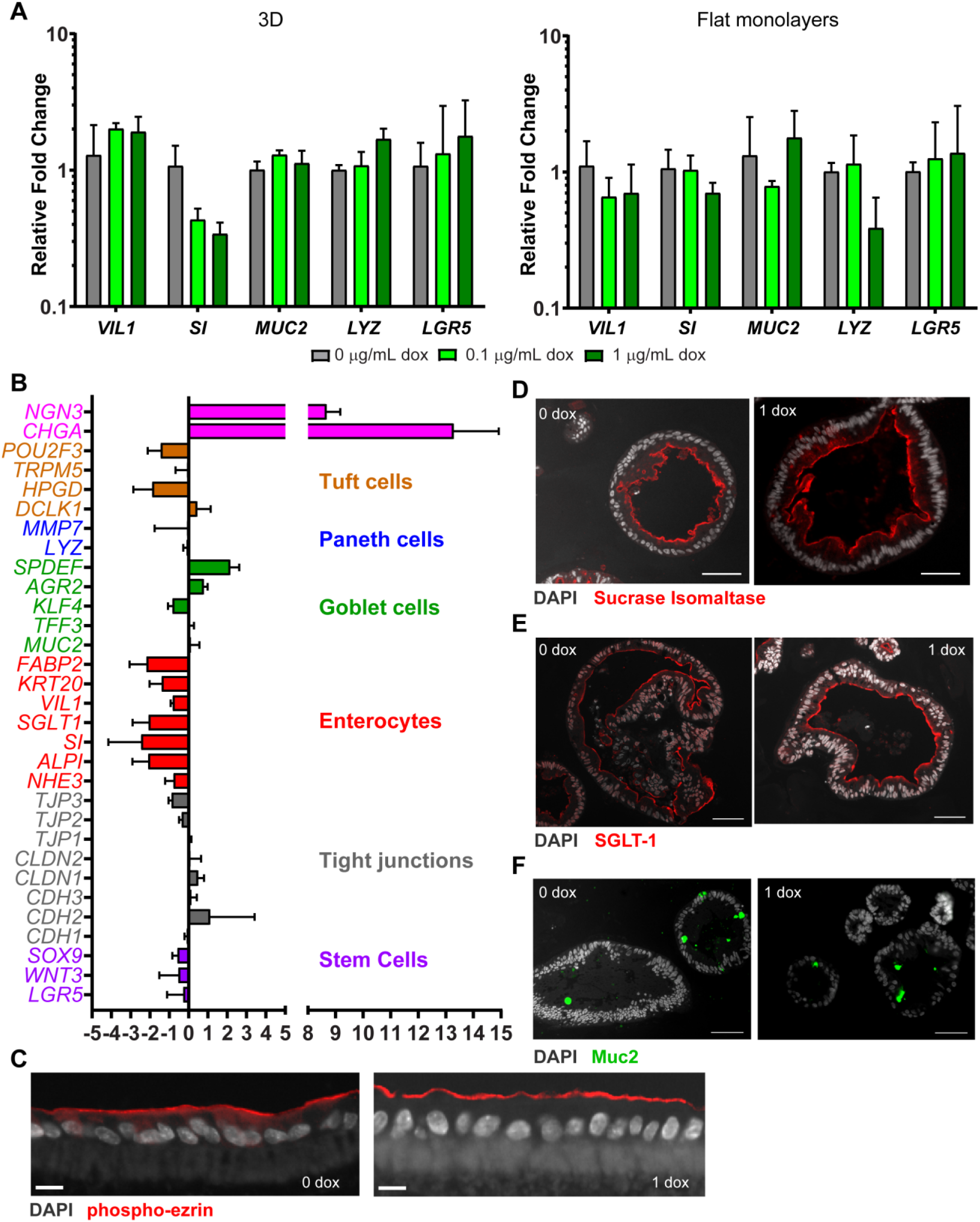
Characterization of epithelial cell types in doxycycline-induced tet*NGN3*-HIEs. **(A)** qPCR of cell marker mRNA transcripts normalized to 18S mRNA transcripts in tet*NGN3*-HIE 3D cultures and flat monolayers (n = 3 biological replicates) **(B)** Log_2_ fold expression of FPKM (fragments per kilobase of transcript per million mapped reads) values from RNA-seq analysis. Transwell monolayers **(C)** and 3D cultures **(D-F)** were treated with 0 or 1 µg/mL doxycycline and fixed and stained. IF staining for phospho-ezrin (Alexa Fluor 555 [red]) **(C)**, sucrase isomaltase (Alexa Fluor 555 [red]) **(D)**, and sodium-glucose transporter 1 (Alexa Fluor 555 [red]) **(E)** to mark the apical border and **(F)** for muc2 (Alexa Fluor 488 [green]) to mark goblet cells and counterstained with DAPI (blue). Images were acquired using a 20X Plan Apo (NA 0.75) DIC objective on an upright Nikon Eclipse 90i microscope with a Photometrics CoolSNAP HQ2 camera and Nikon Elements software. (**C** scale bar = 10µm, **D-F** scale bar = 50µm)

To gain further insight into the broader impacts of *NGN3* overexpression, we performed global transcriptional analysis of mRNA (RNA-seq) isolated from tet*NGN3-* HIEs cultured in transwell format in the absence or presence of 1 µg/mL doxycycline for 5 days. Comparison of the expression levels across 31 genes indicated a dramatic increase in the abundance of *NGN3* and *CHGA* transcripts (**Fig. 4B**) However, there was relatively little change in genes involved in tight junctions, or markers of Paneth, goblet or tuft cell lineages. In contrast, all markers of enterocytes decreased modestly, with log_2_ fold changes of less than 2 (**Fig. 4B, Table 4**). Similar to the H&E staining (**Fig. 3**), immunofluorescence microscopy of 3D cultures and transwell monolayers demonstrated that the tet*NGN3*-HIEs maintained a polarized cell layer upon doxycycline treatment, as demonstrated by apical localization of phospho-ezrin, SI, and sodium-glucose transporter 1 (**Fig. 4C-E**). Finally, we examined whether *NGN3* overexpression altered goblet cell numbers in the tet*NGN3*-HIEs. Immunostaining for Muc2 in non-induced or induced (1 µg/mL doxycycline) 3D tet*NGN3*-HIEs showed that induction of *NGN3* expression did not decrease the number of Muc2-positive cells (**Fig. 4F**), which is consistent with the lack of changes in *MUC2* gene expression between 0 and 1 µg/mL doxycycline treatments found by qPCR and RNA-seq analyses (**Fig. 4A,B**). Taken together, inducible *NGN3* overexpression significantly increases the EEC population, but this does not substantially change the transcript levels of other differentiated cell types or the morphological characteristics of these enteroids.

**Table 4:** RNA-seq Analysis of Fold Changes in Gene Expression with Doxycycline Treatment

| Gene Symbol | Gene Name | Marker of | Fold change FPKM | SD | Log<sub>2</sub> fold change FPKM | SD |
| --- | --- | --- | --- | --- | --- | --- |
| *LGR5* | Leucine-rich repeat containing G protein coupled receptor 5 | Stem cells | 0.94 | 0.50 | -0.28 | 0.842 |
| *WNT3a* | Wnt family member 3A | Stem cells | 0.81 | 0.39 | -0.53 | 0.993 |
| *SOX9* | SRY-Box 9 | Stem cells | 0.68 | 0.12 | -0.58 | 0.267 |
| *CDH1* | Cadherin 1 | Tight Junctions | 0.93 | 0.08 | -0.10 | 0.122 |
| *CDH2* | Cadherin 2 | Tight Junctions | 4.80 | 4.09 | 1.12 | 2.303 |
| *CDH3* | Cadherin 3 | Tight Junctions | 1.12 | 0.23 | 0.14 | 0.285 |
| *CLDN1* | Claudin1 | Tight Junctions | 1.43 | 0.29 | 0.49 | 0.302 |
| *CLDN2* | Claudin 2 | Tight Junctions | 1.14 | 0.47 | 0.09 | 0.552 |
| *TJP1* | Tight junction protein 1 | Tight Junctions | 1.03 | 0.08 | 0.03 | 0.114 |
| *TJP2* | Tight junction protein 2 | Tight Junctions | 0.79 | 0.08 | -0.35 | 0.143 |
| *TJP3* | Tight junction protein 3 | Tight Junctions | 0.55 | 0.07 | -0.88 | 0.159 |
| *NHE3* | Solute carrier family 9 member A3 | Enterocytes | 0.61 | 0.19 | -0.78 | 0.443 |
| *ALPI* | Alkaline phosphatase | Enterocytes | 0.27 | 0.13 | -2.09 | 0.818 |
| *SI* | Sucrose isomaltase | Enterocytes | 0.29 | 0.25 | -2.46 | 1.697 |
| *SGLT* | Solute carrier family 5 member 1 | Enterocytes | 0.27 | 0.14 | -2.07 | 0.837 |
| *VIL1* | Villin | Enterocytes | 0.57 | 0.04 | -0.82 | 0.106 |
| *KRT20* | Keratin 20 | Enterocytes | 0.41 | 0.17 | -1.39 | 0.635 |
| *FABP2* | Fatty acid binding protein 2 | Enterocytes | 0.25 | 0.14 | -2.18 | 0.882 |
| *MUC2* | Mucin 2 | Goblet cells | 1.13 | 0.35 | 0.12 | 0.445 |
| *TFF3* | Trefoil factor 3 | Goblet cells | 1.04 | 0.17 | 0.04 | 0.239 |
| *KLF4* | Kruppel-like factor 4 | Goblet cells | 0.57 | 0.09 | -0.83 | 0.238 |
| *AGR2* | Anterior gradient 2 | Goblet cells | 1.74 | 0.24 | 0.79 | 0.196 |
| *SPDEF* | SAM pointed domain containing ETS transcription factor | Goblet cells | 4.66 | 1.38 | 2.16 | 0.462 |
| *LYZ* | Lysozyme | Paneth cells | 0.94 | 0.12 | -0.10 | 0.178 |
| *MMP7* | Matrix metalloproteinase 7 | Paneth cells | 1.53 | 1.46 | -0.08 | 1.676 |
| *DCLK1* | Double cortin like kinase 1 | Tuft cells | 1.52 | 0.82 | 0.45 | 0.687 |
| *HPGD* | 15-hydroxyprostaglandin dehydrogenase | Tuft cells | 0.33 | 0.21 | -1.88 | 0.981 |
| *TRPM5* | Transient receptor potential cation channel subfamily M member 5 | Tuft cells | 1.01 | 0.40 | -0.08 | 0.593 |
| *POU2F3* | POU class 2 homeobox 3 | Tuft cells | 0.40 | 0.19 | -1.45 | 0.675 |
| *NGN3* | Neurogenin-3 | Enteroendocrine | 428.67 | 141.38 | 8.68 | 0.494 |
| *CHGA* | Chromogranin A | Enteroendocrine | 17779.46 | 23032.47 | 13.29 | 1.63 |
| *TPH1* | Tryptophan hydroxylase 1 | Enteroendocrine | 3054.77 | 251.17 | 11.17 | 1.25 |
| *PAX6* | Paired Box 6 | Enteroendocrine | 75.60 | 43.61 | 5.97 | 0.96 |
| *NEUROD1* | Neuronal differentiation 1 | Enteroendocrine | 5073.90 | 2738.32 | 11.83 | 1.39 |
| *GIP* | Gastric inhibitory polypeptide | Enteroendocrine – subtype K | 6926.92 | 3346.00 | 11.26 | 2.63 |
| *GHRL* | Ghrelin prepropeptide | Enteroendocrine<br>– subtype K | 1576.15 | 1351.91 | 10.33 | 0.90 |
| *SST* | Somatostatin | Enteroendocrine<br>– subtype D | 6733.26 | 3495.26 | 10.93 | 2.62 |
| *PAX4* | Paired Box 4 | Enteroendocrine<br>– subtype D | 3468.87 | 1504.07 | 11.32 | 1.22 |
| *GLP1* | Glucagon-like peptide 1 | Enteroendocrine<br>– subtype L | 15.62 | 11.63 | 3.54 | 1.13 |
| *PYY* | Peptide YY | Enteroendocrine<br>– subtype L | 5.65 | 2.68 | 1.73 | 0.646 |
| *GAST* | Gastrin | Enteroendocrine<br>– subtype G | 46.49 | 35.50 | 4.53 | 2.04 |
| *SCT* | Secretin | Enteroendocrine<br>– subtype S | 7.37 | 3.617 | 2.301 | 1.47 |
| *CCK* | Cholecystokinin | Enteroendocrine<br>– subtype I | 39.66 | 26.75 | 4.57 | 1.88 |
| *MLN* | Motilin | Enteroendocrine<br>– subtype M | 146280.01 | 22926.82 | 13.55 | 4.13 |

### tetNGN3-HIEs secrete serotonin in response to physiological stimuli

The enterochromaffin cell subtype is an important source of serotonin-mediated signaling, which has multiple effects on intestinal homeostasis, and disruptions in serotonin signaling have been shown to contribute to several GI disorders [44-50]. Serotonin is synthesized via the conversion of L-tryptophan by the enzyme tryptophan hydroxylase (Tph1), and stored in secretory vesicles for stimulus-driven secretion [51]. Therefore, we tested if induction of *NGN3* overexpression in HIEs increased serotonin response to biological stimuli *in vitro*. We confirmed that both *CHGA* and *TPH1* gene expression are upregulated in doxycycline-induced tet*NGN3*-HIEs (**Fig. 5A**). Further, we showed that doxycycline induction (1 µg/mL) increased the number of ChgA and serotonin double-positive cells by fluorescence of 3D tet*NGN3*-HIEs (**Fig. 5B**). These indicate an increase in the number of serotonin-secreting enterochromaffin cells.

**Figure 5:**
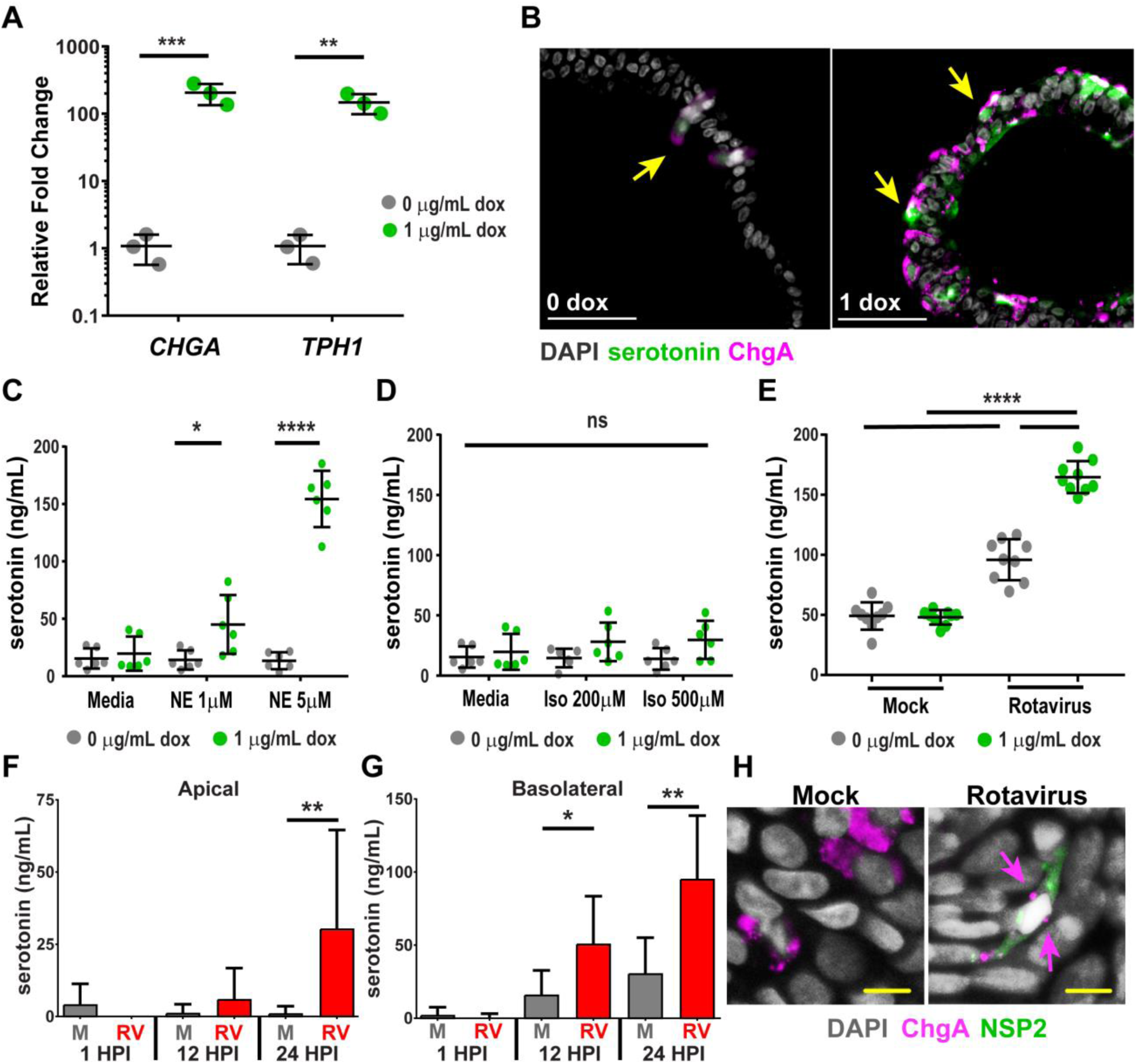
tet*NGN3*-HIEs produce serotonin in response to physiological stimuli. **(A)** qPCR of chromogranin A (*CHGA*) and tryptophan hydroxylase-1 (*TPH1*) mRNA transcripts normalized to 18S mRNA transcripts. (n = 3 biological replicates) **(B)** 3D tetNGN3-HIEs were treated with 0 or 1 µg/mL doxycycline (dox) and fixed and immunostained for chromogranin A (ChgA, Alexa Fluor 555 [pink]) and serotonin (Alexa Fluor 488 [green]) and counterstained with DAPI (gray). Some cells are double-positive for ChgA and seorotnin (yellow arrows). Images were acquired using a 20X Plan Apo (NA 0.75) DIC objective on an upright Nikon Eclipse 90i microscope with a Photometrics CoolSNAP HQ2 camera and Nikon Elements software (scale bar = 50 µm) **(C-E)** tet*NGN3*-HIE flat monolayers were induced with 0 or 1 µg/mL do). Serotonin release as measured by ELISA: **(C)** after 2 hr treatment with norepinephrine (NE) (n = 2 biological replicates), **(D)** after 2 hr treatment with isovalerate (Iso) (n = 2 biological replicates), and **(E)** after 24 hpi with rotavirus (n = 3 biological replicates) **(F-G)** tet*NGN3*-HIE transwell monolayers were differentiated for 5 days with 0.1 µg/mL doxycycline and mock or rotavirus (RV)-infected. Serotonin release measured by ELISA from apical **(F)** or basal **(G)** transwell compartments. **(H)** Max intensity projection images of confocal Z-stack of transwell monolayers fixed and stained for rotavirus nonstructural protein 2 (NSP2) (Alexa Fluor 488 [green]) and ChgA (Alexa Fluor 568 [pink], arrows) and counterstained with DAPI (gray) using a Plan Apo VC 60x Oil DIC objective on a Nikon A1plus point scanning confocal microscope. (scale bar = 10µm; *p<0.05, **p<0.01,***p<0.001, ****p<0.0001).

We next characterized the physiological response of the tet*NGN3-*HIE-derived enterochromaffin cells to stimuli previously shown to elicit serotonin secretion by other enterochromaffin model systems [52, 53]. A previous study found that mouse enterochromaffin cells secrete serotonin in response to norepinephrine and isovalerate [52]. Norepinephrine is a neurotransmitter important for communication between the gut and the enteric nervous system, particularly in response to infection or injury [54]. Isovalerate is a fatty acid metabolite likely generated by the microbiome, particularly by amino acid fermenting bacteria such as from the Clostridial, Bacillus, Lactobacillus, Streptococcus and Proteobacteria groups [55]. We quantified serotonin secretion into the media of flat monolayers of tet*NGN3*-HIEs, induced with 0 or 1 µg/mL doxycycline, and treated with different concentrations of norepinephrine or isovalerate. After stimulation with these agonists, supernatants were collected to measure serotonin secretion using a serotonin ELISA. We found that norepinephrine treatment of induced tet*NGN3*-HIEs stimulated a significant and dose-dependent increase in serotonin secretion, but no increase in serotonin secretion was observed from the non-induced tet*NGN3*-HIEs (p<0.05, p<0.0001) (**Fig. 5C**). Interestingly, treatment with up to 500 µM isovalerate did not stimulate measurable serotonin secretion in either the non-induced or the doxycycline-induced tet*NGN3*-HIEs (**Fig. 5D**).

To determine if our model is responsive to other microbial stimuli, we tested whether rotavirus (RV) infection, a common diarrhea-causing enteric virus, stimulates serotonin secretion from tet*NGN3*-HIEs. Previous studies in both human enterochromaffin cell lines, and in mice found that RV infects both enterocytes and ChgA- and serotonin-positive enterochromaffin cells and stimulates serotonin secretion [12, 53]. Furthermore, HIEs are a new model for RV infection and enterochromaffin cells in HIEs support RV infection; however, whether RV infection stimulates serotonin secretion in HIEs has not been tested [13, 14]. Flat monolayers of tet*NGN3*-HIE induced with 0 or 1 µg/mL doxycycline were infected with trypsin-activated human RV (strain Ito, G3[P8]). Supernatants were harvested at 24 hr post-infection to measure serotonin secretion. RV infection significantly increased serotonin secretion in both the non-induced and doxycycline-induced tet*NGN3*-HIEs compared to mock-infected tet*NGN3*-HIEs (p< 0.0001) (**Fig. 5E**). Most notably, the serotonin response to RV infection was significantly greater from the doxycycline-induced tet*NGN3*-HIEs, demonstrating that the tet*NGN3*-HIEs amplify observed enterochromaffin cell responses (**Fig. 5E**).

The ability of RV to stimulate serotonin secretion in tet*NGN3*-HIEs was also tested in transwell monolayers that were induced with 0.1 µg/mL doxycycline in differentiation media for 5 days. In transwell monolayers, we were able to test serotonin secretion both apically and basolaterally after infection with RV at 1 hpi, 12 hpi, and 24 hpi (**Fig. 5F and 5G**). At 12 hpi, RV-infected transwell monolayers secreted significantly more serotonin into the basolateral compartment than mock-infected monolayers (p<0.05) (**Fig. 5G**), while apical serotonin secretion was not significantly increased (**Fig. 5F**). By 24 hpi, there was significantly more serotonin secretion both apically and basolaterally from RV-infected than mock-infected monolayers (p<0.01) (**Fig. 5G and 5F**). These data indicate that serotonin is primarily secreted basolaterally from the monolayer, particularly early in RV infection, but both basolateral and apical serotonin secretion is detected later in infection. Finally we confirmed that the doxycycline-induced enterochromaffin cells are susceptible to RV infection by immunofluorescence confocal microscopy. At 12 hpi, the tet*NGN3*-HIE transwell monolayers were fix and stained for ChgA and the RV non-structural protein 2 (NSP2), which shows co-immunostaining of cells for ChgA and NSP2 (**Fig. 5H**) that is consistent with the known susceptibility of enterochromaffin cells to RV infection [12, 13, 53].

### tet NGN3-HIEs differentiate into multiple enteroendocrine cell types

To identify other EEC types that are present in the induced tet*NGN3*-HIEs, we examined the RNA-seq data from doxycycline-induced te*tNGN3*-HIEs for expression of marker genes predominantly expressed in the seven EEC subtypes, as well as marker genes for enterochromaffin cells. We found that expression of each of the EEC marker genes was upregulated to varying degrees (**Fig. 6A, Table 4**), suggesting that doxycycline treatment of the tet*NGN3*-HIEs could induce the other EEC cell types in addition to enterochromaffin cells. These results are supported by positive immunostaining for GLP-1 distinct from ChgA-positive cells in tet*NGN3*-HIE monolayers when induced with 1 µg/mL doxycycline (**Fig. 6B**). To test if doxycycline-induced tet*NGN3*-HIEs functionally amplified the physiological response of these EEC subtypes, we quantified gut hormones known to be secreted by different EEC types after stimulation with norepinephrine or infection with RV, as in the experiments above (**Fig. 6C-F**). We did not detect secreted hormones from uninduced tet*NGN3*-HIE monolayers after stimulation with norepinephrine or RV infection, with the exception of monocyte chemoattractant protein-1 (MCP-1), which is also known to be produced by enterocytes (**Fig. 6C and 6D**). In contrast, tet*NGN3*-HIE monolayers induced with 1 µg/mL doxycycline exhibited increased secretion of MCP-1 and glucose-dependent insulinotropic peptide (GIP, from K cells) in response to 5 µM norepinephrine (**Fig. 6B**). Of note, norepinephrine did not stimulate PYY (L cells) or ghrelin (P/D1 cells). Further, in response to RV infection, the induced tet*NGN3*-HIEs secreted large quantities of MCP-1 and GIP and moderate amounts of PYY and ghrelin (**Fig. 6E**), and this response was absent in uninduced tet*NGN3* HIEs (**Fig. 6D**). Norepinephrine and RV infection did not stimulate PP (pancreatic polypeptide; PP cells also known as F cells) secretion, which is primarily produced in pancreatic cells and only rarely in the intestine [56]. Together, these data show that our tet*NGN3*-HIE model drives the differentiation of multiple functional EEC subtypes that secrete gut hormones in response to biologically relevant stimuli.

**Figure 6:**
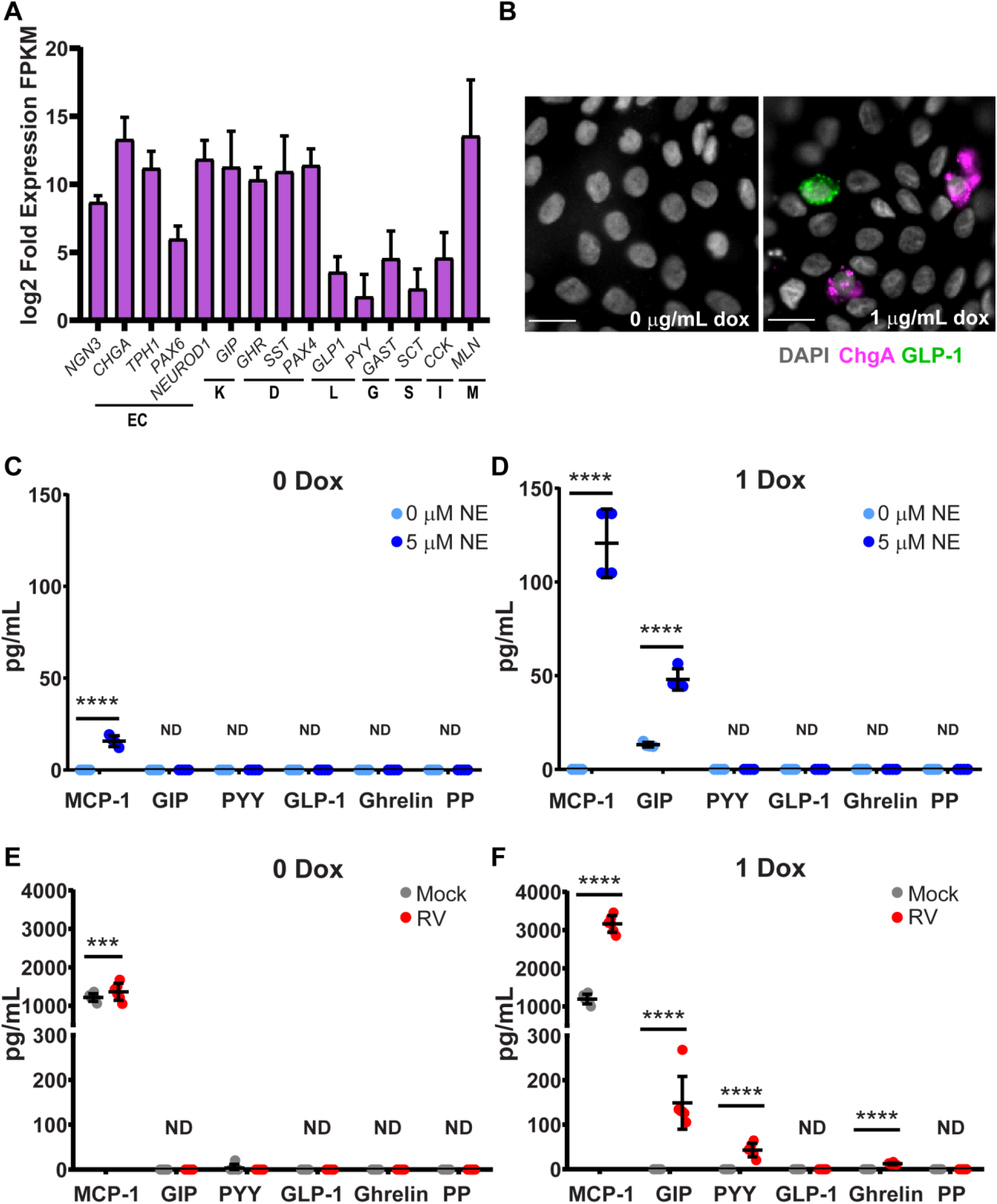
tet*NGN3*-HIEs differentiate into other EEC types. **(A)** log_2_ fold expression of FPKM (fragments per kilobase of transcript per million mapped reads) values from RNA-seq. **(B)** Images of monolayers fixed and stained for chromogranin A (ChgA) (Alexa Fluor 488 [pink]) and glucagon-like peptide-1 (GLP-1) (Alexa Fluor 568 [green]) counterstained with DAPI (gray). Images were acquired using a 40X Apo (NA 1.15) DIC water objective on an inverted Nikon TiE microscope with an ORCA-Flash 4.0 sCMOS camera and Nikon Elements software. Supernatants from tet*NGN3*-HIEs were induced with 0 µg/mL doxycycline (dox) **(C,E)** or 1 µg/mL dox **(D,F)** and collected after **(C,D)** 2 hr stimulation with norepinephrine (NE) (n = 2 biological replicates) or **(E,F)** 24 hpi with rotavirus (RV) (n = 3 biological replicates) and secreted products quantitated by Luminex [monocyte chemoattractant protein-1 (MCP-1), glucose-dependent insulinotropic peptide (GIP), peptide YY (PYY), glucagon-like peptide-1 (GLP-1), pancreatic polypeptide (PP)]. (scale bar = 20µm, ***p<0.001, ****p<0.0001)

## Discussion

Limitations of the available human EEC systems have made it challenging to comprehensively study the molecular physiology of EECs. In this study, we engineered a HIE line to have inducible overexpression of *NGN3* that enable vast upregulation of EEC numbers in the HIE. For the characterization of the tet*NGN3*-HIEs, we initially focused on enterochromaffin cells because they are the dominant intestinal EEC subtype and of great interest for EEC biology, and serve a key role in sensing/responding to microbial metabolites and intestinal pathophysiology [57-59]. We demonstrated that doxycycline-induced *NGN3* overexpression in human jejunual intestinal enteroids increased EECs, as measured by mRNA and protein expression of the endocrine marker *CHGA*. This response was doxycycline-dose dependent, demonstrating that the tet*NGN3*-HIEs allow for “tunable” customization of the number of EECs in different HIE formats (e.g., 3D, flat monolayers, and transwell monolayers). Further, in response to known stimuli, differentiated tet*NGN3*-HIEs release serotonin and several gut hormones characteristic of K cell, L cell, and enterochromaffin cell populations in a single culture system [12, 52, 53].

The tet*NGN3*-HIE model system described in this work overcomes some of the limitations associated with the existing models of EECs. In contrast to primary EECs isolated from tissue and cancer-derived cell lines, HIOs and HIEs are more robust and renewable model systems for studying EECs [29, 52, 60, 61]. However, like the intestine, few EECs spontaneously develop in both HIOs and HIEs, so the rarity of EEC cells remains a limitation of native organoid/enteroid cultures. Thus, increasing *NGN3* expression to drive EEC differentiation, as has been done in recent work with HIOs, is an effective strategy to increase the abundance of EECs for study [29, 40, 62]. In HIOs, *NGN3* overexpression increased the number of ChgA-positive cells to 5% using adenovirus-based constitutive expression and to 23% using lentivirus-based doxycycline-inducible expression [29, 40, 62]. This is comparable to our results with the tet*NGN3*-HIEs wherein we obtained 40% ChgA-positive cells using a lentivirus-based inducible *NGN3* overexpression strategy. These results indicate that for both HIOs and HIEs, stably engineering the stem cell to have inducible expression of *NGN3* is an efficient approach to drive differentiation of higher numbers of EECs.

Together, the use of inducible *NGN3* expression in HIOs and HIEs establish complementary systems that will be widely useful to interrogate different aspects of EEC interactions in the GI tract. In the tet*NGN3*-HIEs model, the HIEs are derived from undifferentiated adult stem cells, and will therefore be particularly useful for studies of adult intestinal epithelial cell expression, function, and physiology. Indeed, isolation, establishment, and transduction of enteroids from individual patients to assess EEC function or response to drug treatments may be a future strategy for personalized medicine [63]. Whereas HIOs are composed of both epithelial and mesenchymal cells, HIEs consist of only epithelial cells so the tet*NGN3*-HIEs allow direct study of EEC interactions with other intestinal epithelial cells. Additionally, co-culture studies that combine the tet*NGN3*-HIEs with other cell types, such as immune cells, endothelial cells, and neurons, will be facilitated by the adaptability of HIEs to different culture formats (3D, 2D, and transwells), which is unique to HIE cultures. Finally, induction and differentiation of tet*NGN3*-HIEs yields EECs in 6-7 days, which may be a simpler and faster system for generating increased numbers of EECs for higher-throughput studies of EEC responses to microbial, diet, or environmental stimuli.

The induced EECs from tet*NGN3*-HIEs exhibit strong functional responses to both norepinephrine and RV infection. Both norepinephrine and RV infection increased secretion of serotonin, MCP-1, and GIP, but RV infection also elicited secretion of PYY and ghrelin. Thus, this system displays distinct functional responses to different stimuli. HIEs have been established as a new, biologically relevant system to study RV-induced gut hormone secretion. RV infects enterochromaffin cells in HIEs and induces serotonin secretion both in mice and *in vitro* from GOT1 cells [12-14]. Here we show that RV infection of tet*NGN3*-HIEs induced serotonin secretion, but this response was significantly amplified by doxycycline-induction to increase EECs. Using transwell monolayers, we were able to show that serotonin is secreted both apically and basolaterally, though it is possible that some basolateral-to-apical diffusion of serotonin occurs, potentially through damaged tight junctions [12, 53]. RV also induced secretion of GIP, PYY, and ghrelin, which was only detected from doxycycline-induced tet*NGN3*-HIEs and indicates that biological responses from rare EECs may be missed in native enteroids. These finding highlights the potential tet*NGN3*-HIEs have for molecular discovery, because secretion of GIP, PYY, and ghrelin during RV infection has not previously been identified, so whether RV infects these EEC subtypes or secretion of these hormones occurs through other signaling pathways merits future investigation [64]. Finally, since insults to the epithelium elevate levels of catecholamines (e.g., norepinephrine), the sensitivity of tet*NGN3*-HIEs to norepinephrine makes them a good model for co-culture studies to measure EEC responses to inflammation or infection [48, 87].

The tet*NGN3*-HIEs are a robust and versatile system for induction of EECs, but some limitations and unknowns about this new model remain. First, the induction of EECs in this system requires lentivirus-integration of a recombinant *NGN3* gene, making these an engineered intestinal organoid system. An alternative approach to induce endogenous EECs in mouse intestinal organoids was established through the combined inhibition of WNT, Notch, and EGFR (i.e., MAPK) signaling pathways [65]. This method leads to increased EEC differentiation, with a concomitant decrease in Paneth and goblet cell numbers [65]. In contrast, the tet*NGN3*-HIEs maintained markers of both Paneth and goblet cells, suggesting that Ngn3 expression alone is insufficient to abolish these other lineage pathways. Second, using this HIE model, we observed increases in most EEC subtypes by RNA-seq and measured hormone secretion from enterochromaffin, L, K, and P/D1 cell subtypes. Although beyond the scope of this study, further studies are needed to determine whether the other subtypes are also functionally responsive to stimuli [66-71]. Finally, the tet*NGN3*-HIEs did not secrete serotonin in response to the short-chain fatty acid isovalerate, in contrast to recent studies using enteroids derived from ChgA-EGFP reporter mice (segment not specified) [52]. Inherent differences between the two systems, such as different receptor expression between human and mouse or between intestinal segments, may account for the different responses.

The tet*NGN3*-HIE system is a new and powerful tool for investigating the roles of EECs in microbial-environment-host communication, infection, inflammation, metabolism. Doxycycline-induction of the tet*NGN3*-HIEs generates large numbers of EECs and amplifies EEC responses to biologically relevant stimuli, yet these cultures retain the salient characteristics of HIEs. Further, this is an adaptable system of in terms of both regulating the number of EECs and culture formats, which enables greater throughput and functional characterization of EEC responses to a multitude of stimuli. The ability of tet*NGN3*-HIEs to differentially respond to norepinephrine and RV infection suggests this new model system is well-suited to study the EEC modulatory effects of commensal microbes, microbial metabolites, pathogens and inflammatory mediators involved in epithelial health and disease.

## Materials and Methods

### Establishment of HIE cultures

Human intestinal enteroid (HIE) cultures were generated from crypts isolated from the jejunal tissues of adult patients undergoing bariatric surgery as previously described [13]. These established cultures were obtained at Baylor College of Medicine through the Texas Medical Center Digestive Diseases Center Study Design and Clinical Research Core. Three-dimensional HIE cultures were prepared from the tissue samples and maintained in culture as described previously [13, 17]. For these studies, jejunum HIEs from patient J2 were used. Complete medium without growth factors (CMGF-) and complete medium with growth factors (CMGF+) were prepared as previously described [13, 17]. Briefly, CMGF-consisted of Advanced Dulbecco’s modified Eagle medium (DMEM)/F-12 medium (Invitrogen) supplemented with 100 U/mL penicillin-streptomycin (Invitrogen), 10 mM HEPES buffer (Invitrogen), and 1X GlutaMAX (Invitrogen). CMGF+ consisted of CMGF-medium with 50% (v/v) Wnt3A-conditioned medium, 20% (v/v) R-spondin conditioned medium, 10% (v/v) Noggin-conditioned medium, 1X B-27 supplement (Invitrogen), 1X N-2 supplement (Invitrogen), 1 mM N-acetylcysteine (Sigma-Aldrich), 50 ng/mL mouse epidermal growth factor (EGF) (Invitrogen), 10 mM nicotinamide (Sigma-Aldrich, St. Louis, MO), 10 nM Leu-Gastrin I (Sigma-Aldrich), 500 nM A-83-01 (Tocris Bioscience), and 10 nM SB202190 (Sigma-Aldrich).

Differentiation medium consisted of the same components as CMGF+ without Wnt3A conditioned medium, R-spondin conditioned medium, SB202190, and nicotinamide and only 5% (v/v) Noggin conditioned medium. High Wnt CMGF+ (hW-CMGF+), for creating and maintaining lentivirus-transduced HIEs, consisted of CMGF+ mixed with an additional 50% (v/v) Wnt3a conditioned medium. All HIEs were passaged in phenol red-free, growth factor-reduced Matrigel (Corning).

### Creation of NGN3-expressing HIEs

Tetracycline-inducible tet*NGN3*-expressing HIEs were created using lentivirus transduction. The tet*NGN3* lentivirus transfer plasmid was in the pINDUCER backbone and a kind gift from Dr. Noah Shroyer and Dr. Jim Wells [62, 72]. The tet*NGN3* lentiviruses were packaged and harvested in-house with the packaging plasmid psPAX2, a gift from Dr. Didier Trono (Addgene plasmid #12260), and the envelope plasmid pCMV-VSV-G, a gift from Dr. Bob Weinberg (Addgene plasmid #8454), as described in more detail previously [73]. Virus packaging was assessed using Lenti-X GoStix (Clontech) but virus titers were not otherwise measured. Densely seeded jejunum HIEs were grown in hW-CMGF+ for one week to increase stem cell counts. HIEs were dissociated from Matrigel with ice-cold 1X phosphate buffered saline (PBS) and pipetted into 1.7 mL centrifugation tubes for centrifugation in a swinging bucket rotor at 200 x *g* for 5 min at 4°C. Supernatant was removed, HIEs were resuspended and washed twice with 1 mL ice-cold 1X PBS and centrifuged using the same conditions. Supernatants were removed and HIEs resuspended with lentivirus inoculum consisting of 8 µg/mL polybrene (EMD Millipore), 10 µM Y-27632 Rock inhibitor (Tocris Biosciences), 50 µL tet*NGN3* lentivirus, and freshly made hW-CMGF+ for a total volume of 200 µL. HIEs were incubated with the inoculum in closed centrifugation tubes for 24 hr at 37°C in a humidified 5% CO_2_ incubator. After 24 hr, HIEs were centrifuged again and washed twice with ice-cold 1X PBS followed by suspension in Matrigel drops on a 24-well plate. Following Matrigel polymerization in the incubator for 10 min, 500 µL of hW-CMGF+ medium was added to the well. Media was changed every other day for one week before passaging and selecting with 200 µg/mL Geneticin (VWR). We refer to this transduced line as tet*NGN3*-HIEs, and they were grown in hW-CMGF+ with 200 µg/mL Geneticin.

### Preparation and differentiation of HIE monolayers

HIE monolayers were prepared from three-dimensional cultures and seeded into flat 96-well plates or transwells as described previously [21, 42]. In brief, 96-wells or transwell inserts (Costar, cat. no. 3413) were pre-treated with Matrigel diluted in 1XPBS (1:40) and incubated at 37°C. 3D HIEs were lifted from Matrigel and washed with an ice cold solution of 0.5 mM EDTA in 1X PBS and dissociated with 0.05% trypsin/0.5 mM EDTA for 4 min at 37°C. Trypsin was inactivated with CMGF- + 10% FBS and the cell solution was pipetted vigorously and filtered using a 40 µm nylon cell strainer (Falcon, cat. no. 352340) to dissociate into single cells. Then cells were centrifuged for 5 min at 400 x *g*, resuspended with CMGF+ and 10 µM Y-27632 Rock inhibitor, and plated into prepared wells. After 48 hr in CMGF+ and 10 µM Y-27632 Rock inhibitor, the medium was changed to differentiation media with the addition of 10 µM Y-27632 Rock inhibitor and indicated concentrations of doxycycline (Fisher Scientific). Differentiation medium with Y-27632 and doxycycline was changed every day for 4-5 days to differentiate cells.

### Immunofluorescence of HIE monolayers

Two-dimensional HIE flat monolayers were fixed using the BD Cytofix/Cytoperm kit (BD Biosciences, cat. no. 554714) according to manufacturer instructions. Primary antibodies were diluted in BD Perm/Wash buffer and (**Table 1**) were incubated at 4°C overnight. Primary antibodies were recognized by the appropriate secondary antibodies (**Table 1**) and incubated for 1 hr at room temperature. Nuclei were stained with DAPI (Thermo Fisher Scientific, cat. no. R37606) for 5 min at room temperature and washed with 1X PBS for imaging and storage.

**Table 1:** Antibodies used in this work

| Antibody Type | Target | Species | Company | Cat# | Dilution |
| --- | --- | --- | --- | --- | --- |
| Primary | Muc2 | Mouse | Santa Cruz | sc-515032 | 1:200 |
| Primary | Sucrase Isomaltase | Mouse | Santa Cruz | sc-393470 | 1:100 |
| Primary | Sodium-glucose transporter 1/ SLC5A1 | Rabbit | Novus Biologicals | NBP-238748 | 1:100 |
| Primary | Chromogranin A | Rabbit | Novus Biologicals | NBP-253140 | 1:500 |
| Primary | Serotonin | Goat | Immunostar | 20079 | 1:500 |
| Primary | Phospho-Ezrin | Rabbit | Cell Signaling | 3726S | 1:200 |
| Primary | Rotavirus NSP2 | Guinea Pig | [74] |  | 1:500 |
| Secondary | Anti-goat Alexa Fluor 488 | Donkey | Life Technologies | A11055 | 1:1,000 |
| Secondary | Anti-rabbit Alexa Fluor 488 | Donkey | Life Technologies | R37116 | 1:2,000 |
| Secondary | Anti-rabbit Alexa Fluor 555 | Donkey | Life Technologies | A31572 | 1:1,000 |
| Secondary | Anti-guinea pig Alexa Fluor 568 | Goat | Life Technologies | A11075 | 1:1,000 |
| Secondary | Anti-mouse Alexa Fluor 568 | Goat | Life Technologies | A11004 | 1:2,000 |

### Immunofluorescence of paraffin-embedded sections

Three-dimensional (3D) HIEs embedded in Matrigel were gently removed and transferred to 1 mL syringes. The cells were then fixed in 4% (v/v) paraformaldehyde solution for 1 hr at room temperature. After fixation, cells were deposited into a cassette for paraffin embedding. For transwell cross-sections HIE membranes were cut from transwells, fixed in 4% (v/v) paraformaldehyde solution for 1 hr at room temperature and placed into cassettes for paraffin embedding. Paraffin-embedded sections of 7 µm in thickness were subjected to a series of dehydration steps. Epitope retrieval was performed by incubating slides with Vector Labs Antigen Unmasking Solution Citrate Buffer pH 6 (Vector labs, cat no. H-3300) for 20 min at 100°C in a steamer. Slides were then blocked for 1 hr at room temperature in 10% goat and/or donkey serum. Primary antibodies (**Table 1**) were incubated at 4°C overnight. Primary antibodies were recognized by the appropriate secondary antibodies (**Table 1**) and incubated for 1 hr at room temperature. Nuclei were stained with DAPI (Thermo Fisher Scientific, cat. no. R37606) for 5 min at room temperature. All slides were cover-slipped with mounting media (Life Technologies) and imaged.

### Transmission Electron Microscopy

Three-dimensional HIEs embedded in Matrigel were primary fixed in a solution of 2% paraformaldehyde + 2.5% glutaraldehyde + 2 mM calcium chloride in 0.1 M cacodylate buffer (pH = 7.4) for 5-7 days at 4°C. They were post-fixed in 1% osmium tetroxide in 0.1 M cacodylate buffer for 1 hr and en bloc stained with saturated aqueous uranyl acetate. After a routine dehydration sequence, tissue pieces were gradually infiltrated in a gradient series of Spurr’s Low Viscosity resin and ethanol, then embedded in fresh Spurr’s resin and polymerized at 60° C for 3 days. 55-60 nm thin sections were cut on a Leica UC7 ultra-microtome using a Diatome Ultra 45 diamond knife. Sections were viewed on a Hitachi H7500 transmission electron microscope set to 80kV. Images were collected using an AMT XR-16 digital camera and AMT Image Capture, v602.600.51 software.

### Microscopy and image analysis

tet*NGN3*-HIEs were imaged with widefield epifluorescence on a Nikon TiE inverted microscope and an upright Nikon Eclipse 90i microscope using a SPECTRA X LED light source (Lumencor) as well as a Nikon A1plus point scanning confocal microscope for fluorescence imaging. The following objectives were used: 10X Plan Fluor (NA 0.3) phase contrast objective, 20X Plan Apo (NA 0.75) differential interference contrast (DIC) objective, a 40X Apo DIC water objective, and a Plan Apo VC 60X Oil DIC objective. Fluorescence images were recorded using either an ORCA-Flash 4.0 sCMOS camera (Hamamatsu) or a CoolSNAP HQ2 camera (Photometrics), and color images for H&E sections were recorded using a DS-Fi1-U2 camera (Nikon). Nikon Elements Advanced Research v4.5 software was used for data acquisition and image analysis.

To quantify ChgA-positive cells in 3D and monolayer tet*NGN3*-HIEs, images were analyzed using Nikon Elements software. Individual images were processed with a 4 µm size threshold by channel to reduce noise from nonspecific staining. Images were morphologically separated using a 3×3 matrix, and objects touching the borders were removed. The percent of positive cells was the number of Alexa Fluor 488-positive objects divided by the number of DAPI-stained detected nuclei. At least 5 images per condition were analyzed with an average of 140 (3D cultures) or 1400 (flat monolayers) nuclei per image.

### RNA extraction, reverse transcription, and real-time PCR

tet*NGN3*-HIEs were rinsed once with ice cold 1XPBS and transferred to a centrifugation tube (technical duplicates were combined into a single tube). Cells were lysed by the addition of 1 mL of TRIZOL Reagent (Invitrogen) and mixed thoroughly using a vortex mixer for 30 seconds. Chloroform (200 µL) was added, the samples were mixed again, and then incubated for 5 min at room temperature. The phases were separated by centrifugation at 14,000 × *g* for 10 min, and the aqueous phase was moved to a new tube (∼400 µL). Total RNA was then isolated using the RNeasy Isolation Kit (Qiagen) according to the manufacturer’s instructions. RNA was eluted in 20 µL sterile nuclease free water (Fischer Scientific). RNA concentration and purity were determined by absorbance at 260 nm and 280 nm using a spectrophotometer (DS-11, DeNovix). RNA (200 ng) was treated with DNase (Ambion Turbo DNA-free) to remove any contaminating genomic DNA per manufacturer’s directions, and then cDNA was synthesized from purified total RNA using SuperScript III reverse transcriptase (Invitrogen) according to the manufacturer’s instructions. Reaction components included random hexamers (Integrated DNA Technologies), 10 mM dNTPs (New England Biosciences), and approximately 2 ng of RNA. No reverse transcriptase controls were included by omitting SuperScript III.

Real time PCR reactions were performed in triplicate with the following reaction components: 1 µL cDNA, 0.25 µL of 20 mM forward and reverse primer (**Table 2**), 10 µL Power SYBR Green PCR MasterMix (Applied Biosystems), and 8.5 µL nuclease free water (Fisher Scientific). Real-time PCR was performed using QuantStudio real time thermocycler (Applied Biosciences) under the following conditions: 95°C 10 min, 40 cycles of 95°C for 15 seconds, followed by 60°C for 1 min, when data was recorded. A melting curve between 60°C and 95°C was performed. No template controls were included by substituting water for the cDNA in the reaction. C_T_ values of technical replicates were averaged for each independent biological experiment and normalized to the expression of the 18S ribosomal subunit. Data graphed are representative of 3 experiments (n=3 biological replicates, with 3 technical replicates in each experiment).

### Preparation of HIE transwells for RNA-sequencing and transcriptional analysis

To assess the transcriptional profile changes of tet*NGN3-HIEs* during doxycycline induction. HIE monolayers were prepared on transwells and differentiated as described above. The total RNA was extracted from transwells in the presence or absence of 1 µg/ml doxycycline using the same protocol as for RT and qPCR described above. rRNA integrity was checked on an Agilent 2100 bioanalyzer from six independent biological replicates per condition. Paired-end Illumina sequencing was performed on mRNA enriched samples by Novogene (USA) following the standard work-flow procedure. Raw sequence reads were mapped to human genome hg19 using STAR software to generate the number of fragments per kilobase per million mapped reads (FPKM) for each gene. Ratios of transcript abundance per group were based on the log_2_ FPKM value in 1 µg/ml doxycycline condition relative to the 0 µg/mL doxycycline condition were used to determine the fold change in gene expression. Samples were filtered for those containing >10% N value, presence of adaptor sequences, and quality. Greater than 97% of reads in each sample were clean reads post-filtering with error rates ≤ 0.03%. Pearson correlation coefficients between samples were between 0.92 and 0.97 R^2^ values. In total, 10,741 genes were differentially expressed between the groups (5,870 up-regulated, 4,871 down-regulated) as indicated by DESeq2 analysis using a padj value <0.05, and a log_2_ fold change >1, q-value <0.005.

### Isovalerate and norepinephrine stimulation

After four days in differentiation medium and induced with 0, 0.1, or 1 µg/mL doxycycline, confluent HIE monolayers were washed with 1XPBS. Isovalerate (Sigma-Aldrich) and noradrenaline bitartrate (Tocris Bioscience) solutions were prepared in 1X PBS and added to the HIE monolayers. Monolayers were incubated for 2 hr at 37°C and 5% CO_2_. Following incubation, supernatants were harvested and kept at −20°C for downstream analysis. PrestoBlue Cell Viability Reagent (ThermoFisher Scientific), a resazurin-based assay, was utilized according to manufacturer’s recommended protocols. Fluorescence intensity of reduced resazurin (560-610 nm, Infinite F200Pro, Tecan) measures metabolic activity of the cells and was used as an indication of cell viability.

### Rotavirus infection of HIE monolayers

African green monkey kidney (MA104) cells were cultured in DMEM supplemented with 10% (v/v) fetal bovine serum (Corning). Ito RV (G3[P8]) was propagated in MA104 cells in serum-free DMEM in the presence of 1 µg/mL Worthington’s Trypsin (Worthington Biochemical). After harvest, stocks were subjected to three freeze/thaw cycles. Plaque assays with MA104 cells were used to determine the titers of the viral preparations. The tet*NGN3*-HIE flat or transwell monolayers were induced with 0, 0.1, or 1 µg/mL doxycycline in differentiation media for 4-5 days. For infection, the confluent HIE monolayers were washed once with CMGF-. Mock MA104 cell lysates and Ito RV were treated with 10 µg/mL Worthington’s trypsin for 30 min at 37°C. Then the tet*NGN3*-HIE monolayers were treated with an inoculum of CMGF- with MA104 cell lysate or Ito RV. Cells were infected basolaterally following observations that this method results in more efficient infections (Blutt, Crawford, and Estes, unpublished data). Transwell and flat monolayers were incubated for 1 hr or 2 hr, respectively, at 37°C with 5% CO_2_. Following incubation, the supernatants were removed and replaced with differentiation media and returned to the incubator. For the 1 hpi timepoint, supernatants were collected 5 min after washing inoculum. Supernatants were harvested and kept at −20°C for downstream analyses.

### Measurement of serotonin release

Serotonin secretion by HIEs following stimulation with norepinephrine, isovalerate, or RV infection was quantified by ELISA (Eagle Biosciences) according to the manufacturer’s instructions. A standard curve of known serotonin concentrations was plotted against optical density at 450 nm with a limit of detection of 2.6 ng/mL (Infinite F200Pro, Tecan).

### Measurement of metabolic hormones

Hormone secretion was quantified by Luminex assay (Human Metabolic Hormone Magnetic Bead Panel, HMHEMAG-34K, EMD Millipore) according to manufacturer’s standard protocol. Supernatants following stimulation with norepinephrine or RV infection were assayed for the amount of secreted amylin (active), C-Peptide, ghrelin (active), GIP (total), GLP-1 (total), glucagon, IL-6, leptin, MCP-1, PP, PYY (total), and TNF-α. Limits of detection of the assay are: amylin 11.81 pg/ml, C-Peptide 13.28 pg/ml, ghrelin 9.40 pg/ml, GIP 0.33 pg/ml, GLP-1 1.11 pg/ml, glucagon 12.73 pg/ml, IL-6 3.75 pg/ml, leptin 54.16 pg/ml, MCP-1 8.57 pg/ml, PP 0.68 pg/ml, PYY 11.18 pg/ml, and TNF-α <0.14 pg/ml.

### Statistical analysis

Biostatistical analyses were performed using GraphPad Prism (version 7) software (GraphPad Inc., La Jolla, CA). Comparisons were made with either One-way or Two-way Analysis of Variance (ANOVA) and Tukey’s post hoc multiple comparisons test when appropriate. Differences between the groups were considered significant at p < 0.05 (*), and the data are presented as mean ± standard deviation. All authors had access to the study data, reviewed, and approved the final manuscript.

## Author Contributions

Concept and design (ACG, HAD, MAE, CTD, UCK, MKE, RAB, JMH); intellectual contribution (ACG, HAD, MAE, CTD, UCK, MKE, JV, RAB, JMH); data acquisition (ACG, HAD, MAE, CTD, UCK); data analysis, statistical analysis, and interpretation (ACG, HAD, MAE, CTD, UCK); drafting and editing manuscript (ACG, HAD, MAE, JMH); obtained funding (MKE, JV, RAB, JMH)

## Abbreviations

(NGN3): neurogenin-3
(tet): tetracycline
(dox): doxycycline
(GI): gastrointestinal
(HIEs): human intestinal enteroids
(HIOs): human intestinal organoids
(SCFAs): short chain fatty acids
(ChgA): chromogranin A
(PP): pancreatic polypeptide
(GIP): glucose-dependent insulinotropic peptide
(GLP-1): glucagon-like peptide-1
(PYY): peptide YY
(MCP-1): monocyte chemoattractant protein-1
(Tph1): tryptophan hydroxylase
(VIL1): villin
(SI): sucrase isomaltase
(RV): rotavirus
(hr): hour
(hpi): hours post infection
(RT-qPCR): reverse transcriptase – quantitative polymerase chain reaction
(IF): immunofluorescence
(H&E): hematoxylin and eosin
(2D): two-dimensional
(3D): three-dimensional
(CMGF-): complete medium without growth factors
(CMGF+): complete media with growth factors
(hW-CMGF+): high Wnt complete media with growth factors

## Acknowledgments

We would like to thank Dr. Noah Shroyer for providing the tet*NGN3* plasmid. The BCM Medical Scientist Training Program and the Integrative Molecular and Biomedical Sciences Graduate Program provided additional trainee support (A.C.G). We would like to thank Xi-Lei (Shelly) Zeng and Xiaomin Yu of the Digestive Diseases Center Enteroid Core for their help with maintaining the enteroid culturing and enteroid media. We thank Debra Townley and Michael A. Mancini, Ph.D. of the Integrated Microscopy Core for their help with transmission electron microscopy of the enteroids.

## Graphical Abstract

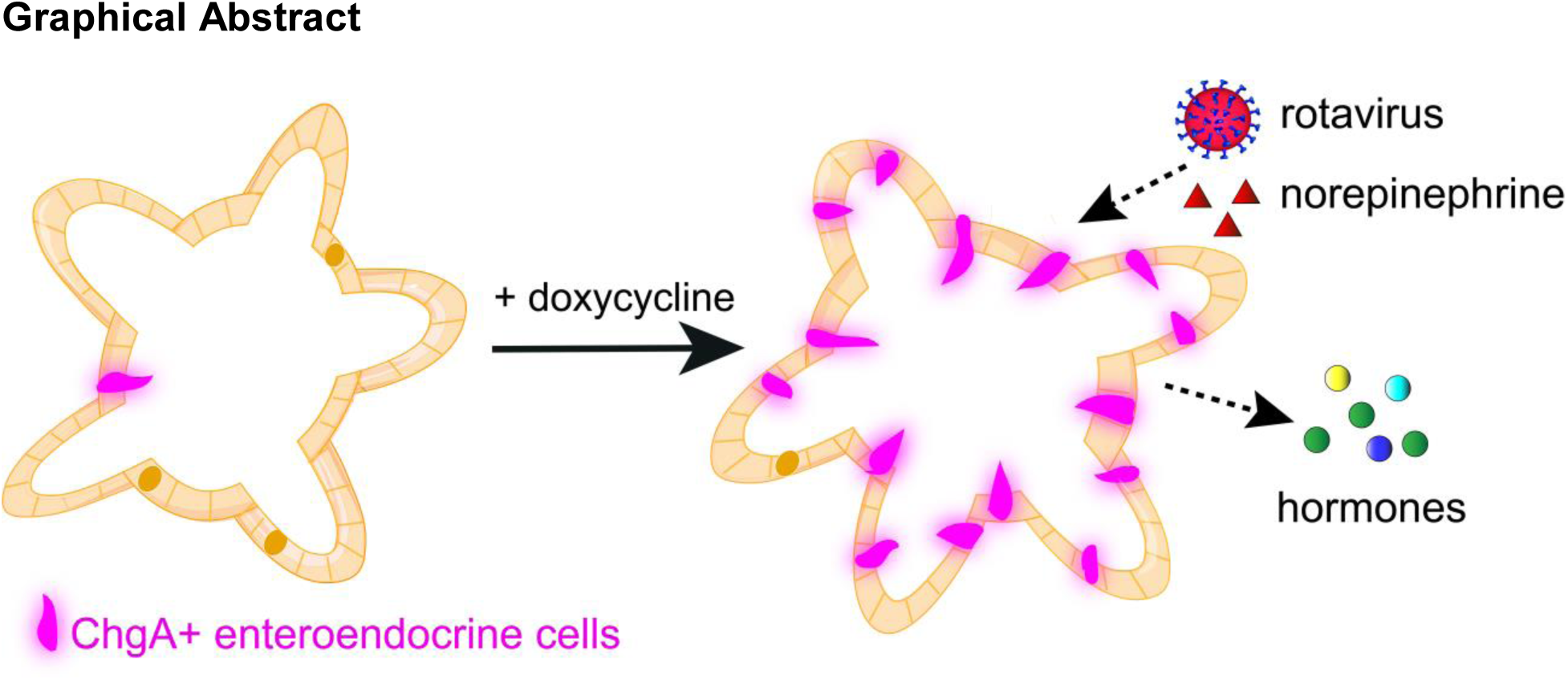

